# Symptomatic treatment by a BBB-permeable AAV engineered to restore TDP-43 function slows motor neuron disease and prevents paralysis

**DOI:** 10.1101/2025.08.14.670400

**Authors:** Aswathy Peethambaran Mallika, Jessica G. Yu, Opal Sitzman, Meghraj Singh Baghel, Shruti Renganathan, Irika R. Sinha, Tatiana Melnikova, Jonathan P. Ling, Philip C. Wong

## Abstract

TAR DNA-binding protein 43kDa (TDP-43) dysfunction is an early pathogenic mechanism that underlies amyotrophic lateral sclerosis (ALS), a devastating neurodegenerative disorder that lacks disease modifying therapies. We previously developed a mouse model in which TDP-43 is selectively deleted from motor neurons (*ChAT-Cre;Tardbp^f/f^*) that mimics the early stages of ALS. Here, we demonstrate that intravenous delivery of a blood-brain-barrier (BBB) permeable AAV capsid expressing our rationally designed splicing repressor CTR (AAV-PHP.eB-CTR) in symptomatic *ChAT-Cre;Tardbp^f/f^* mice markedly slowed disease progression and prevented paralysis. Systemic delivery of AAV-PHP.eB-CTR led to transduction of ∼80% of spinal motor neurons, repression of TDP-43-associated cryptic exons within motor neurons expressing CTR, and attenuation of motor neuron loss. Notably, the addition of the *TARDBP* 3’UTR autoregulatory element to CTR maintained its expression within a physiological range. In control littermates that received AAV-PHP.eB-CTR and were monitored for >20 months, grip strength and body weight remained normal, and no histopathological abnormalities were observed, underscoring a favorable safety profile for this gene therapy. These results provide preclinical proof-of-concept that BBB-crossing AAV delivery of CTR can rescue motor neuron disease through the restoration of TDP-43 function, offering a promising mechanism-based therapeutic strategy for ALS.

## INTRODUCTION

Emerging evidence indicates that deficits associated with TAR DNA-binding protein 43 (TDP-43) cryptic splicing (Irwin et al., 2024; Seddighi et al., 2024; Brown et al., 2022; Ma et al., 2022; Klim et al., 2019; Melamed et al., 2019; Ling et al., 2015) are a central pathogenic mechanism underlying Amyotrophic Lateral Sclerosis-Frontotemporal Dementia (ALS-FTD). TDP-43, an RNA-binding protein, was first linked to ALS-FTD through the observation of hallmark cytoplasmic inclusions within patient neurons (Neumann et al., 2006). A key function of TDP-43 is to repress cryptic exons, intronic sequences whose inclusion disrupts mature mRNA transcripts and contributes to neuronal dysfunction in ALS-FTD (Ling et al., 2015). More recently, deficits in TDP-43 splicing repression have been detected in presymptomatic ALS (Irwin et al., 2024) and shown to precede TDP-43 cytoplasmic aggregation by at least a decade in the aging human brain (Chang et al., 2023), suggesting that loss of TDP-43 nuclear function contributes to neuronal degeneration. This interpretation is strengthened by evidence that ALS-linked TDP-43 mutations (Gitcho et al., 2008; Kabashi et al., 2008; Sreedharan et al., 2008) can promote the inclusion of the *STMN2* cryptic exon in human iPSC-derived neurons, independent of TDP-43 cytoplasmic aggregation (Klim et al., 2019; Melamed et al., 2019; Prudencio et al., 2020). At the genetic level, a major risk allele for ALS-FTD promotes the inclusion of a cryptic exon in *UNC13A* that is normally repressed by TDP-43 (Brown et al., 2022; Ma et al., 2022; van Es et al., 2009). Furthermore, TDP-43 nuclear depletion can occur without cytoplasmic inclusions in certain presymptomatic FTD cases (Nana et al., 2019; Vatsavayai et al., 2016) and in Alzheimer’s disease cases with TDP-43 pathology (AD-TDP) (Calliari et al., 2024; Sun et al., 2017). These findings support a model in which early loss of nuclear TDP-43 function initiates cryptic splicing defects during the presymptomatic stage of disease and can drive neurodegeneration. Thus, therapeutic strategies designed to restore TDP-43 function represent a potential mechanism-based therapy for ALS-FTD.

We previously developed a conditional knockout mouse model lacking TDP-43 in spinal motor neurons (*ChAT-Cre;Tardbp^f/f^* mice), which exhibits motor neuron disease resembling the prodromal phase of ALS (Donde et al., 2019). In this model, we showed that TDP-43 dependent splicing repression is critical for motor neuron survival and developed an AAV-based therapeutic strategy to restore this function using a novel splicing repressor we termed CTR (Donde et al., 2019; Ling et al., 2015). In this study we aimed to model the clinically relevant therapeutic context by evaluating the safety and efficacy of AAV-mediated CTR by treating mice after symptom onset. Accordingly, we delivered the CTR payload to adult *ChAT-Cre;Tardbp^f/f^* mice at the symptomatic stage of disease using the blood-brain barrier (BBB) crossing AAV-PHP.eB serotype via an intravenous (IV) route of administration (Chan et al., 2017; Goertsen et al., 2022). Systemic delivery of AAV-PHP.eB-CTR efficiently transduced motor neurons, restored TDP-43 dependent splicing repression, and mitigated motor neuron disease phenotype. Furthermore, the incorporation of a TDP-43 3’UTR autoregulatory element as a build-in “safety switch” constrained CTR expression to physiological levels in motor neurons and long-term exposure of AAV-PHP.eB-CTR in all aged mice revealed no adverse events. Together, these results support a favorable safety profile and provide critical validation of AAV-mediated CTR delivery as a clinically viable therapeutic strategy for ALS.

## RESULTS

### Intravenous delivery of AAV-PHP.eB-CTR leads to efficient transduction of motor neurons in symptomatic *ChAT-Cre;Tardbp^f/f^* mice

To model TDP-43 pathology in ALS, we utilized our *ChAT-Cre;Tardbp^f/f^* mouse line, in which the *ChAT-Cre* driver selectively deletes TDP-43 in motor neurons (Donde et al., 2019). These mice develop a progressive motor neuron disease that ultimately leads to premature death. Using a previously established breeding scheme, we generated a large cohort of *ChAT-Cre;Tardbp^f/f^* mice and littermate controls (see Methods & **Supplementary Fig. 1**). CTR was delivered by lateral tail vein injection of AAV-PHP.eB-CTR at the optimized dose of 5E+13 vg/Kg (**Supplementary Fig. 2**). Injections were performed at 6 weeks of age, when fine tremors are already apparent in *ChAT-Cre;Tardbp^f/f^* mice, resulting in accumulation of CTR in motor neurons throughout the brain stem and spinal cord. The transduction rate of motor neurons in lumbar spinal cord and brain stem was 79±6.2% and 72±11.7%, respectively, as determined by CTR immunostaining at one-month post-injection (**Fig. 1A, B & Supplementary Fig. 2**).

**Fig. 1:**
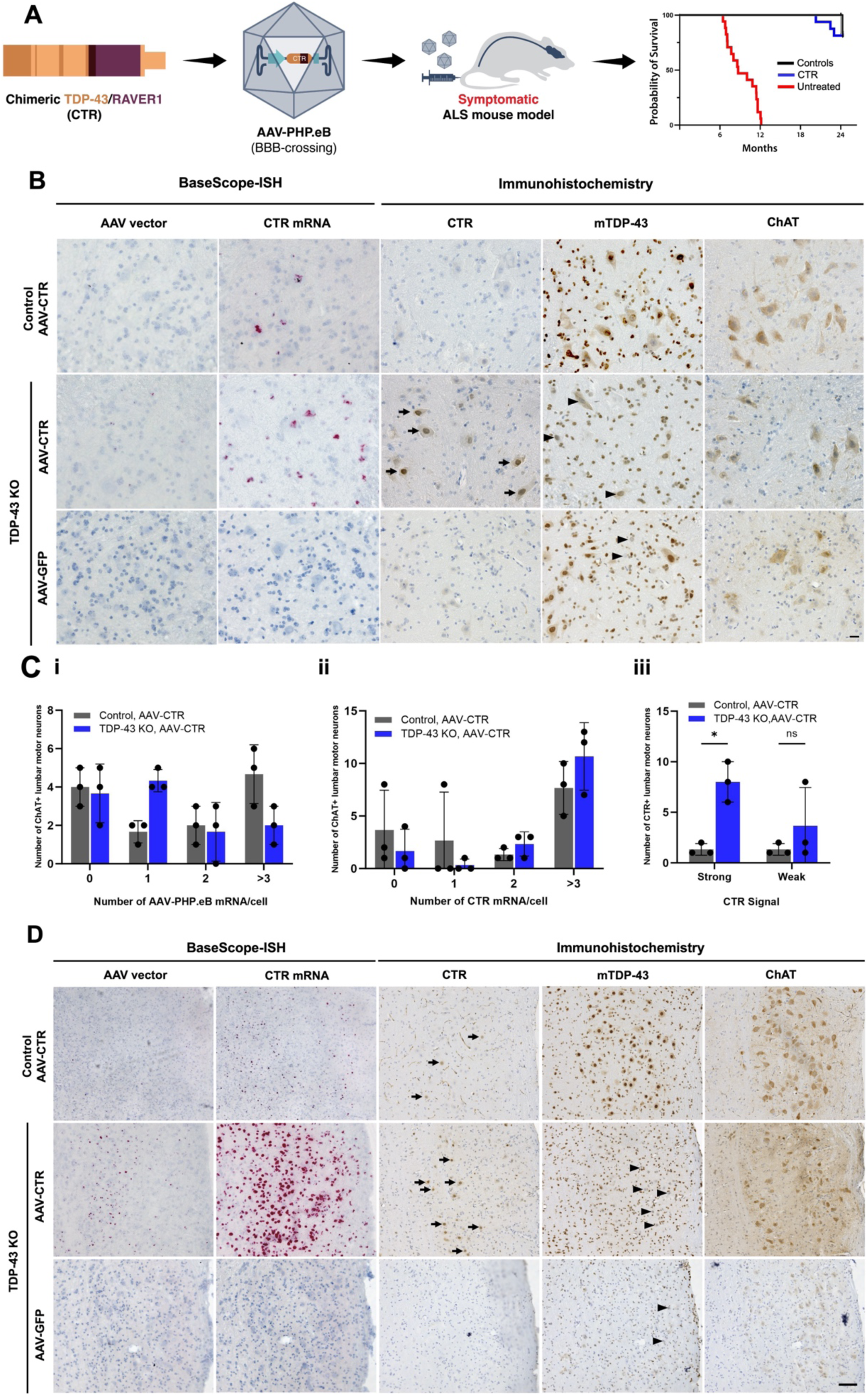
Intravenous delivery of AAV-PHP.eB-CTR confers high efficiency of transduction in adult motor neurons. (**A**) Schematic diagram showing the study design. (**B**) Representative images showing the accumulation of CTR in ventral motor neurons of *ChAT-Cre;Tardbp^f/f^* mice. CTR protein staining was carried out using the anti-human TDP-43 antibody (indicated by arrows in Panel 2). The administration of AAV-PHP.eB-CTR via tail vein injection results in a mean infectivity rate of 79±6.2% (n=3). Immunostaining for mouse TDP-43 indicates ∼60% depletion of endogenous TDP-43 (indicated by arrowheads in Panels 2 and 3) and CTR as detected by anti-human TDP-43 in lumbar spinal cord sections of three- and five-month-old *ChAT-Cre;Tardbp^f/f^* mice. Panel 1. AAV-PHP.eB-CTR treated *ChAT-Cre;Tardbp^f/+^* mice, Panel 2. AAV-PHP.eB-CTR treated *ChAT-Cre;Tardbp^f/f^* mice and Panel 3. AAV-PHP.eB-GFP treated *ChAT-Cre;Tardbp^f/f^* mice. Scale bar=50 µm. (**C**) Single cell quantification of AAV genome copy number (i) and CTR mRNA expression (ii) using BaseScope-ISH probes and immunohistochemical staining of CTR (iii) show that the autoregulatory domain expressed along with CTR prevents the overexpression of CTR in cells with normal expression of TDP-43. (**D**) Representative images showing accumulation of CTR in the facial motor nuclei of *ChAT-Cre;Tardbp^f/f^* mice. The administration of AAV-PHP.eB-CTR via tail vein injection results in a mean infectivity rate of 72±11.7% (n=3). Arrows indicate anti-human TDP-43 immunostaining for CTR, and arrowheads indicate endogenous TDP-43 in Panels 2 and 3. Panel 1. AAV-PHP.eB-CTR treated *ChAT-Cre;Tardbp^f/+^* mice, Panel 2. AAV-PHP.eB-CTR treated *ChAT-Cre;Tardbp^f/f^* mice and Panel 3. AAV-PHP.eB-GFP treated *ChAT-Cre;Tardbp^f/f^* mice. For all groups, n=3. Scale bar=50 µm.

### Autoregulatory domain regulates the level of CTR protein and prevents overexpression toxicity

To prevent the potential toxicity of excessive accumulation of CTR, which would likely mimic the deleterious effects of TDP-43 overexpression (Arnold et al., 2013; Carmen-Orozco et al., 2024; Fratta et al., 2018), we engineered AAV-PHP.eB-CTR to include the endogenous autoregulatory element within TDP-43’s 3’UTR as an intrinsic “safety switch”. We designed probes for BaseScope RED assay, a sensitive RNA *in situ* hybridization method, to detect both the AAV backbone and *CTR* mRNA and to determine the number of *CTR* transcripts within motor neurons of spinal cord and brainstem (**Fig. 1B**). While the genome copy number of AAV-PHP.eB-CTR is maintained at 2-3 copies per cell (**Fig. 1Ci**), the accumulation of *CTR* mRNA in spinal motor neurons was higher in TDP-43 deficient motor neurons of *ChAT-Cre;Tardbp^f/f^* mice when compared to normal motor neurons in *ChAT-Cre;Tardbp^f/+^* mice (**Fig. 1Cii**). These data indicate that *CTR* transcripts are suppressed when TDP-43 is present within a motor neuron. Furthermore, the levels of CTR protein were elevated in spinal motor neurons depleted of TDP-43 as compared to spinal motor neurons in control mice (**Fig. 1Ciii**). Long-term monitoring of control *ChAT-Cre;Tardbp^f/+^* mice treated with AAV-PHP.eB-CTR showed normal grip strength, body weight, and histopathology, with no evidence of vector-related toxicity (**Supplementary Fig. 3**). Together, these findings confirm that the TDP-43 3’UTR autoregulatory element can serve as an effective safety mechanism to prevent over-accumulation of CTR protein in motor neurons.

### Symptomatic delivery of AAV-PHP.eB-CTR to *ChAT-Cre;Tardbp^f/f^* mice significantly slows disease progression and prevents limb paralysis

To determine the impact of AAV-PHP.eB-CTR on motor neuron disease phenotype, we initially assessed the grip strength of *ChAT-Cre;Tardbp^f/f^* mice treated at symptomatic stage. As expected, the GFP control (AAV-PHP.eB-GFP) treated *ChAT-Cre;Tardbp^f/f^* mice showed deficits as early as 6 weeks of age and continued to worsen over the next 6 months (**Fig. 2A**). Remarkably, when compared to control *ChAT-Cre;Tardbp^f/+^* mice, *ChAT-Cre;Tardbp^f/f^* mice treated with AAV-PHP.eB-CTR were able to maintain normal grip strength over this same period of monitoring (**Fig. 2A**). Accompanied by this rescue of grip strength, the body weight of AAV-PHP.eB-CTR treated *ChAT-Cre;Tardbp^f/f^* mice continued to increase, like that of control littermates (**Fig. 2B & Supplementary Fig. 4**). In contrast, the body weight of AAV-PHP.eB-GFP injected *ChAT-Cre;Tardbp^f/f^* mice failed to increase over this same period (**Fig. 2B & Supplementary Fig. 4**). To determine whether improved grip strength and the ability of AAV-PHP.eB-CTR treated *ChAT-Cre;Tardbp^f/f^* mice to maintain normal body weight impacted survival, we monitored the progression of disease until AAV-PHP.eB-GFP treated *ChAT-Cre;Tardbp^f/f^* mice became moribund. While all AAV-PHP.eB-GFP treated *ChAT-Cre;Tardbp^f/f^* mice reached end-stage disease around 47 weeks of age, all AAV-PHP.eB-CTR treated *ChAT-Cre;Tardbp^f/f^* mice remained alive to at least 100 weeks of age (**Fig. 2C**). Among the aged AAV-PHP.eB-CTR treated *ChAT-Cre;Tardbp^f/f^* mice (n=4) or AAV-PHP.eB-CTR or AAV-PHP.eB-GFP treated control littermates (*ChAT-Cre;Tardbp^f/+^*; n=2) that became clinically moribund, three of these animals exhibited marked skin lesions and were subsequently euthanized. Importantly, none of these animals displayed evidence of limb paralysis. These results establish that the BBB permeable AAV-PHP.eB-CTR restores TDP-43 function to motor neurons when delivered intravenously after onset of symptoms in a mouse model that mimics the early phase of ALS. These findings strongly support CTR as a promising AAV gene therapy designed to target TDP-43 dysfunction in ALS.

**Fig. 2:**
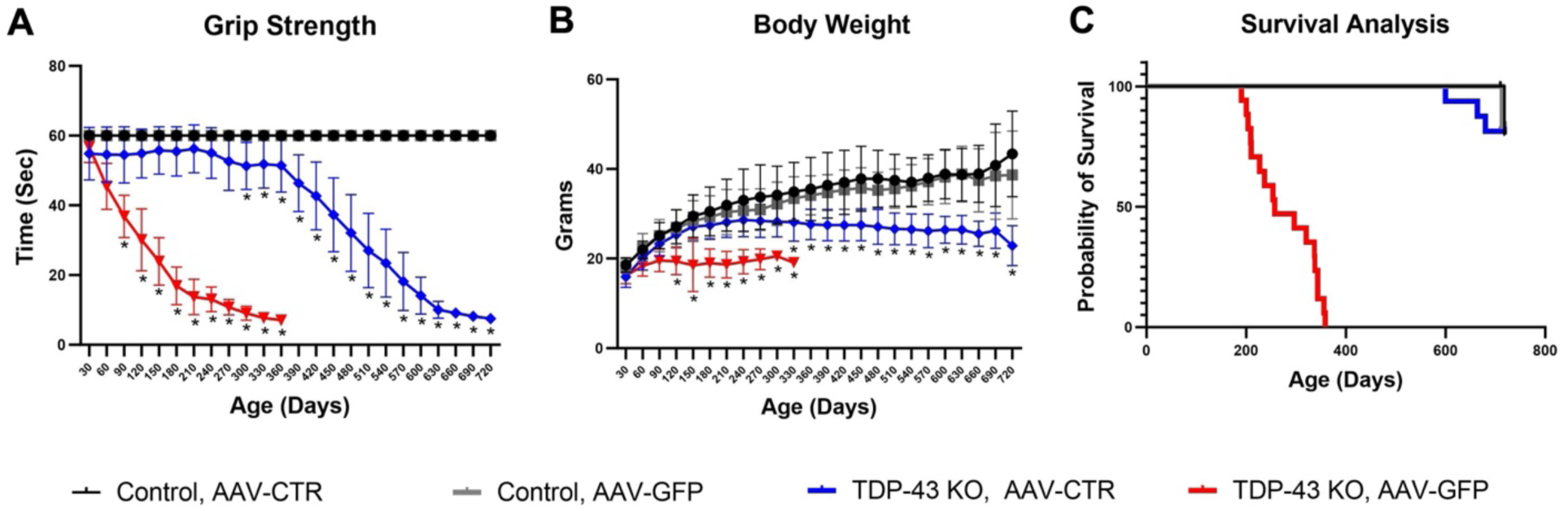
Efficacy of intravenous delivery of AAV-PHP.eB-CTR to *ChAT-Cre;Tardbp^f/f^* mice after symptom onset. (**A**) Grip strength analysis using performance in hanging wire experiments revealed marked amelioration in motor deficits (characteristic of the TDP-43 knockout mice) in CTR treated adult *ChAT-Cre;Tardbp^f/f^* mice (*p<0.0001, two-way ANOVA test). (B) Intravenous administration of CTR resulted in the mitigation of behavioral deficits. Significant improvement was observed in the age-dependent reduction of body weight associated with the *ChAT-Cre;Tardbp^f/f^* mice (*p<0.0001, two way ANOVA using Turkey’s multiple comparison test). (C) Intravenous administration of CTR resulted in the extension of survival. Kaplan-Meier survival curve of *ChAT-Cre;Tardbp^f/+^* and *ChAT-Cre;Tardbp^f/f^* mice administered AAV-PHP.eB expressing either CTR or GFP as control. Data from male and female cohorts aged between 23-24 months are shown together. For all groups, n=15.

### Rescue of motor neuron disease correlates with prevention of cryptic exon inclusion by AAV-PHP.eB-CTR

To determine whether the striking rescue of muscle strength, body weight loss and premature death of *ChAT-Cre;Tardbp^f/f^* mice by AAV-PHP.eB-CTR was correlated with cryptic exon repression and motor neuron survival, we used an integrated co-detection assay (BaseScope RED assay) to monitor both the inclusion of previously characterized cryptic exons such as *Synj2bp* in combination with ChAT or endogenous TDP-43 or CTR immunostaining in motor neurons. Consistent with previous RNA-seq analyses (Brown et al., 2022; Jeong et al., 2017; Klim et al., 2019; Ling et al., 2015; Ma et al., 2022; Melamed et al., 2019), transcripts of these cryptic exons served as molecular markers of TDP-43 loss of function (**Fig. 3A**). We found that CTR prevented inclusion of these cryptic exons in both spinal (**Fig. 3B, D & Supplementary Fig. 5**) and facial (**Fig. 3C, D**) motor neurons of AAV-PHP.eB-CTR treated *ChAT-Cre;Tardbp^f/f^* mice. To determine whether prevention of TDP-43 cryptic exons led to attenuation of motor neuron loss, we quantified the number of ChAT+ motor neurons in 5-month-old *ChAT-Cre;Tardbp^f/f^* mice treated with AAV-PHP.eB-CTR. Using a combination of histological and immunohistochemical techniques, we observed loss of ∼50% of motor neurons in AAV-PHP.eB-GFP-treated *ChAT-Cre;Tardbp^f/f^* mice (**Fig. 4A**). In contrast, AAV-PHP.eB-CTR significantly attenuated the loss of these cells (**Fig. 4B**). Concordant with the rescue of motor neurons in the *ChAT-Cre;Tardbp^f/f^* mice treated with AAV-PHP.eB-CTR, the large diameter fibers of the ventral root were restored (**Fig. 4C, D**) and skeletal muscle atrophy was markedly attenuated (**Fig. 4E & Supplementary Fig. 6**). These results strongly indicate that the rescue of motor neuron disease was conferred by the robust protection of motor neurons due to CTR-mediated repression of TDP-43-associated cryptic exons within ∼80% of motor neurons. Therefore, our findings validate this AAV-based strategy to restore TDP-43 function, offering a promising opportunity to advance this novel mechanism-based gene therapy for the treatment of ALS.

**Fig. 3:**
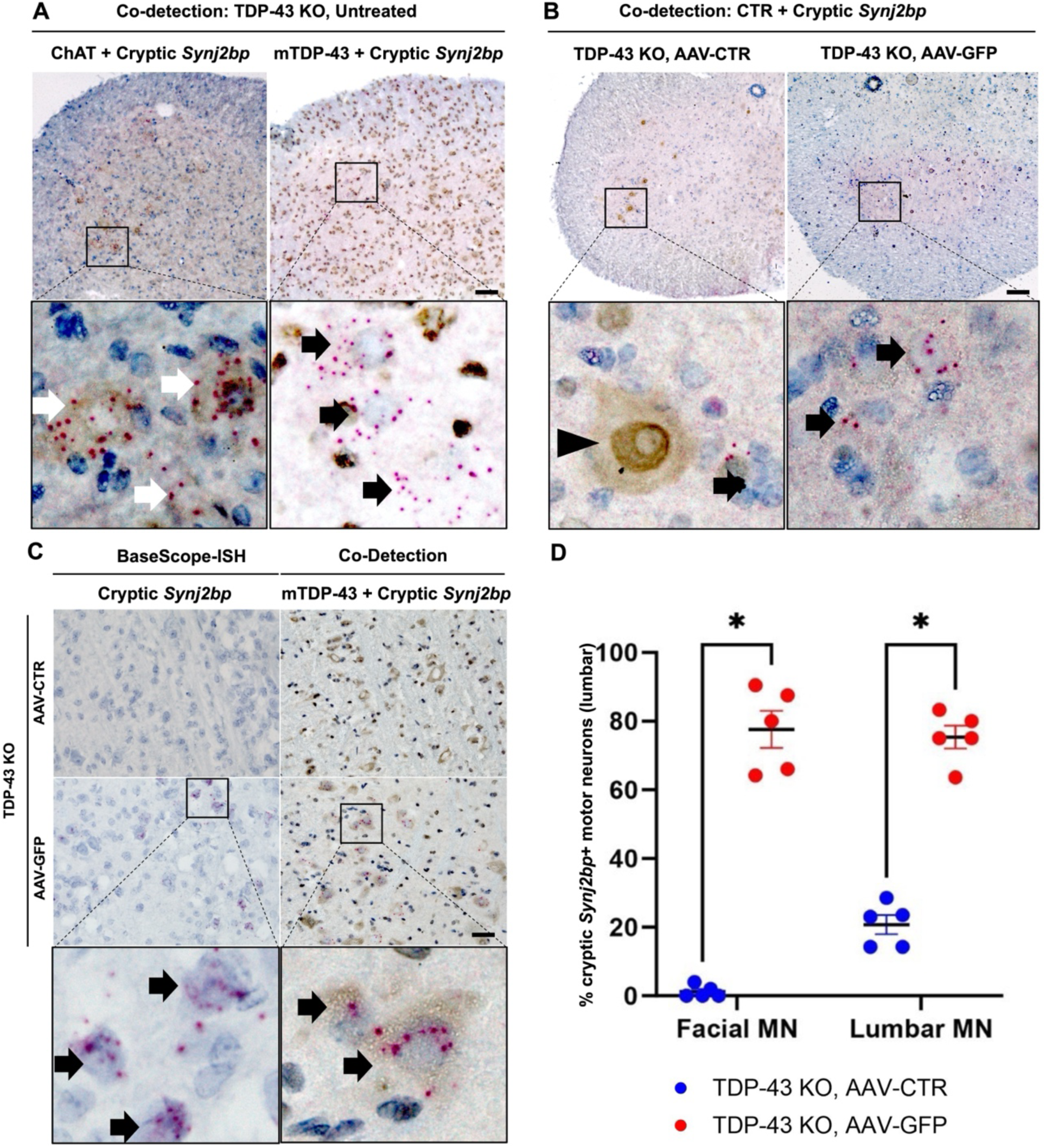
AAV-PHP.eB-CTR prevents inclusion of TDP-43 cryptic exons. (**A**) Representative images of simultaneous detection of the cryptic exon in *Synj2bp* using BaseScope-ISH probe and antibodies against ChAT or mouse TDP-43 reveals an elevated presence of transcripts containing *Synj2bp* cryptic exon in some ChAT+ neurons or in neurons lacking mouse TDP-43 in *ChAT-Cre;Tardbp^f/f^* mice. White arrows indicate *Synj2bp* cryptic exon in ChAT+ neurons. Black arrows indicate *Synj2bp* cryptic exon in TDP-43 deficient cells. Scale bar=100 µm. (**B**) Representative images of co-detection of cryptic exon in *Synj2bp* with CTR (using anti-human TDP-43) shows the rescue of cryptic exon expression in CTR expressing cells in *ChAT-Cre;Tardbp^f/f^* mice. Arrowheads indicate CTR expressing neurons, and black arrows show CTR or mouse TDP-43 deficient cells expressing transcripts containing *Synj2bp* cryptic exon (n=5). Scale bar=100 µm. (**C**) Representative images from BaseScope-ISH or Co-detection assay to detect *Synj2bp* cryptic exon in facial motor nuclei. Arrows depict TDP-43 deficient cells containing transcripts with *Synj2bp* cryptic exons in *ChAT-Cre;Tardbp^f/f^* mice (n=5). Scale bar=50 µm. (**D**) Quantitative analysis of ChAT positive motor neurons expressing *Synj2bp* cryptic exon in *ChAT-Cre;Tardbp^f/f^* mice treated with AAV-PHP.eB-GFP and AAV-PHP.eB-CTR (n=5 per group).

**Fig. 4:**
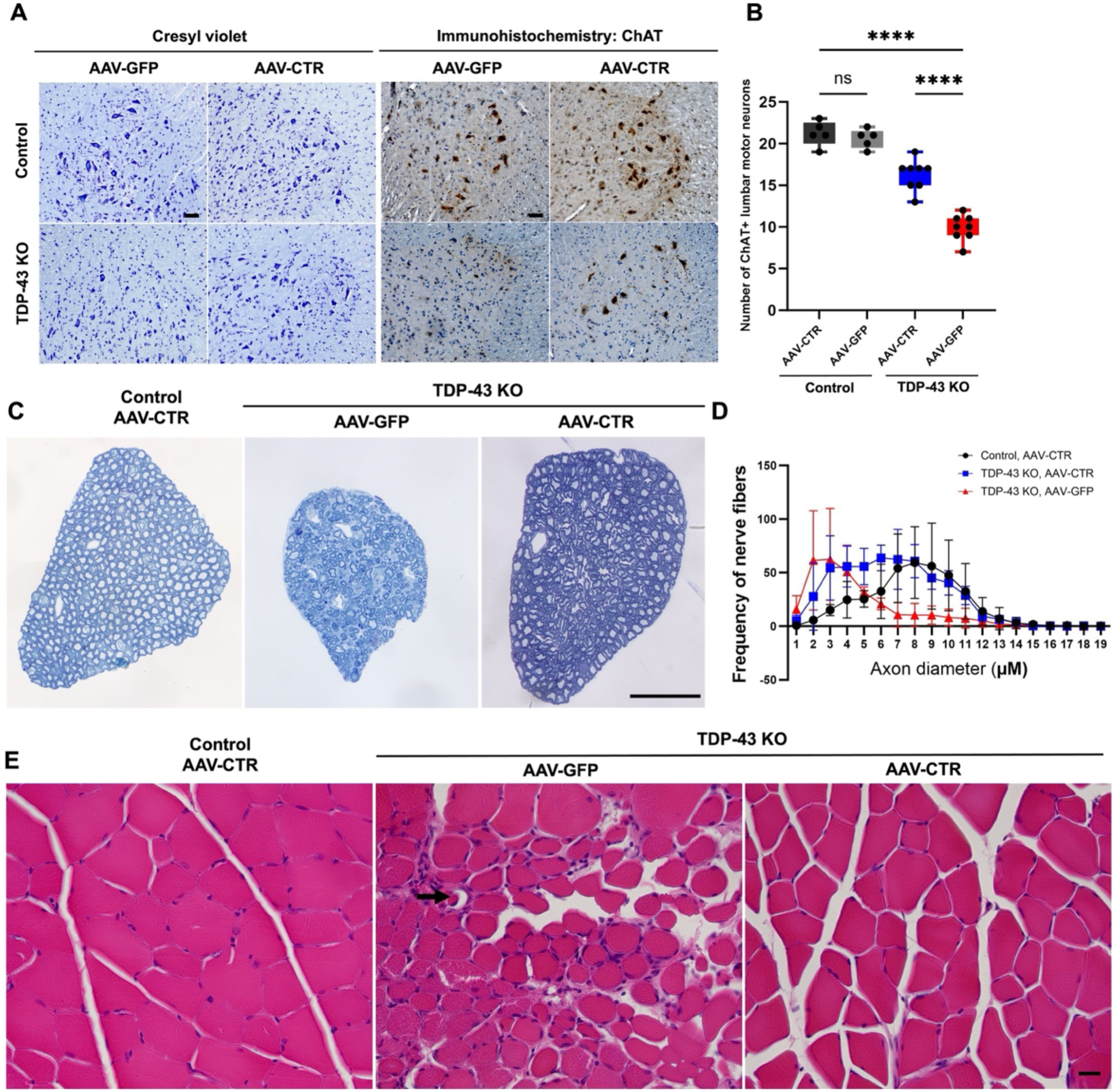
AAV-PHP.eB-CTR attenuates motor neuron loss, axon degeneration, and skeletal muscle atrophy. (**A**) Representative images from pathological analysis showing that CTR protects against motor neuron loss in adult *ChAT-Cre;Tardbp^f/f^* mice treated with AAV-PHP.eB-CTR at symptomatic stage (treated at 1.5 months of age and analyzed at 5 months of age, n=8). (**B**) Quantification of number of ChAT positive motor neurons in the lumbar spinal cord (n=8). Scale bar=50 µm. (**C**) Representative toluidine blue-stained light microscopic images of transversely sectioned motor roots (n=3). (**D**) A notable reduction in the number of large axonal fibers is observed in the knockout mice treated with AAV-PHP.eB-GFP compared to those treated with AAV-PHP.eB-CTR. Scale bar=50 µm. (**E**) Representative images after H&E staining of gastrocnemius muscle sections (10 µm) from *ChAT-Cre;Tardbp^f/f^* treated with AAV-PHP.eB-CTR and *ChAT-Cre;Tardbp^f/f^* mice treated with AAV-PHP.eB-CTR or AAV-PHP.eB-GFP at 5 months of age. Arrows represent the muscle fibers undergoing degeneration in *ChAT-Cre;Tardbp^f/f^* mice treated with AAV-PHP.eB-GFP, which is rescued in *ChAT-Cre;Tardbp^f/f^* mice treated with AAV-PHP.eB-CTR. Scale bar=50 µm.

## DISCUSSION

ALS is a fatal neurodegenerative disease characterized by progressive loss of upper and lower motor neurons, leading to weakness, paralysis, and death within 2-5 years of diagnosis (Taylor et al., 2016). Current approved therapies, such as riluzole and edaravone, offer modest benefits in slowing disease progression or alleviating symptoms (Mead et al., 2023). More recently, an antisense oligonucleotide targeting *SOD1* (tofersen) has demonstrated clinical benefit in patients with *SOD1* mutations (Blair, 2023; Miller et al., 2022). However, *SOD1* mutation carriers represent a small fraction of ALS cases, underscoring the urgent need for additional mechanism-based therapies (Mead et al., 2023; Taylor et al., 2016).

The discovery of nuclear clearance and cytoplasmic inclusions of TDP-43 in cases of ALS and FTD (Neumann et al., 2006), together with the identification of pathogenic missense mutations in the *TARDBP* gene encoding TDP-43 (Kabashi et al., 2008; Sreedharan et al., 2008), has focused efforts toward understanding the disease mechanisms driven by TDP-43 pathology. Because protein aggregation has been a hallmark of many neurodegenerative disorders, early work focused on the potential toxic gain-of-function mechanisms of TDP-43 aggregates (Lee et al., 2011; Taylor et al., 2016). However, it was possible that the loss of normal nuclear TDP-43 function could also play a central role in ALS pathogenesis.

A well-established function of TDP-43 is the repression of cryptic exons during pre-mRNA splicing (Ling et al., 2015). Under normal conditions, cryptic exons are excluded from mature mRNA transcripts, but when TDP-43 function is disrupted, these cryptic exons are spliced into mRNA and frequently introduce premature stop codons that lead to nonsense mediated decay (Jeong et al., 2017; Ling et al., 2015). In some cases, cryptic exon inclusion produces abnormal proteins that may contribute to neurodegeneration or serve as fluid biomarkers (Irwin et al., 2024; Ling et al., 2015). TDP-43 pathology is also linked to broader RNA metabolism defects, including dysregulation of RNA stability, transport, and translation (Balendra et al., 2025; Lagier-Tourenne et al., 2012; Mehta et al., 2023). Among these RNA processing defects, the failure to repress cryptic exons has emerged as a hallmark pathogenic mechanism in ALS (Brown et al., 2022; Klim et al., 2019; Ling et al., 2015; Ma et al., 2022; Melamed et al., 2019; Prudencio et al., 2020; Tan et al., 2016). Our recent work demonstrates that TDP-43 dysregulation occurs early, even at presymptomatic stages of disease (Irwin et al., 2024), providing a strong rationale for therapeutics that restore this loss of function in ALS.

However, prior studies of TDP-43 in transgenic models have demonstrated that increasing the expression of either wildtype or mutant TDP-43 can be independently deleterious, leading to widespread splicing disruptions and neurodegeneration (Arnold et al., 2013; Carmen-Orozco et al., 2024; Fratta et al., 2018). These findings underscore the importance of gene expression regulation for any therapeutic designed to compensate TDP-43 loss-of-function. One mitigation strategy is to regulate transgene levels by incorporating frameshifting cryptic exons whose splicing is dependent on TDP-43 activity (Wilkins et al., 2024; Ling et al., 2022). In our AAV design, we incorporated the endogenous 3’UTR autoregulatory element of TDP-43 to enable CTR to be selectively upregulated only in TDP-43 deficient neurons, which successfully mitigated overexpression toxicity.

In this study, we utilized a model of TDP-43 loss-of-function that captures early ALS pathology (Donde et al., 2019; Ling et al., 2015) to demonstrate that systemic delivery of a BBB-penetrant AAV encoding CTR can restore cryptic exon repression, preserve motor neuron integrity, and maintain neuromuscular function. We evaluated the therapeutic efficacy of this AAV-based approach by administering after symptom onset, modeling a clinically relevant intervention window for ALS. This treatment prevented functional decline and markedly extended survival, with no evidence of limb paralysis even after long term follow-up. In contrast to other therapeutic approaches, which focus on restoring functional gene expression of individual key targets of TDP-43 dysfunction (Baughn et al., 2023; Mehta et al., 2023), our approach aims to restore repression across all cryptic exons, offering a broader therapeutic potential for ALS. Furthermore, emerging next generation AAV serotypes with the potential to cross the human BBB (Huang et al., 2024) will provide a strong foundation for the clinical translation of our mechanism-based gene therapy. Such advances hold promises not only for ALS, but also for other neurodegenerative disorders characterized by TDP-43 proteinopathy, including FTLD-TDP (Neumann et al., 2006), LATE (Nelson et al., 2019), and AD-TDP (Amador-Ortiz et al., 2007; Josephs et al., 2008, 2016; Uryu et al., 2008).

## MATERIALS AND METHODS

### Animals

Our conditional *Tardbp* knockout mice (*Tardbp^f/f^*, Jax Stock 017591) in which exon 3 is flanked by LoxP sites (Chiang et al., 2010) were crossbred with ChAT-dependent Cre driver line (*ChAT-Cre)* transgenic mice on a C57BL/6J background (Jax Stock 006410) to generate a cohort of *ChAT-Cre;Tardbp^f/+^* mice. These heterozygous mice were then interbred to generate *ChAT-Cre/ChAT-Cre;Tardbp^f/+^*, which was subsequently bred with *Tardbp^f/f^* mice to generate a final cohort of *ChAT-Cre;Tardbp^f/+^*(control mice) and *ChAT-Cre;Tardbp^f/f^* (TDP-43 knockout mice) (Supplementary figure 1D). Animals were housed in a 12-hour light/dark cycle with food and water *ad libitum*. All experiments were performed in accordance with the National Institute of Health (NIH) Guidelines and approved by the Animal Care and Use Committee (ACUC) at Johns Hopkins Medicine.

### Experimental Design

The male or female littermates including control and knockout mice from the final breeding cages were randomly divided into four groups; i.e., two groups from each genotype, 1) *ChAT-Cre;Tardbp^f/+^* mice treated with AAV-PHP.eB-GFP (n=20); 2) *ChAT-Cre;Tardbp^f/+^* mice treated with AAV-PHP.eB-CTR (n=20); 3) *ChAT-Cre;Tardbp^f/f^* mice treated with AAV-PHP.eB.GFP (n=20); and 4) *ChAT-Cre;Tardbp^f/f^* mice treated with AAV-PHP.eB-CTR (n=20). For additional analysis, litters from the same breeding pairs were included to analyze the rate of AAV-PHP.eB transduction efficiency at 1-, 3-, and 5-month of age, and to perform cryptic exon and protein expression analysis.

### Viral Vector Packaging

AAV-PHP.eB packaging was performed by Virovek (Hayward, CA). The expression of CTR or GFP was confirmed independently by HeLa cell transduction and protein blot analysis prior to all experiments.

### Intra-Venous Delivery of AAV-PHP.eB

Intravenous delivery of AAV-PHP.eB vector was performed through lateral tail vein injection in 6-week-old mice. Briefly, mice were warmed using an overhead heat lamp for 2-3 minutes to dilate the veins. The animal was lightly anesthetized and restrained using a commercial device (Tail veiner Restrainer, Braintree Scientific, Inc.). The AAV-PHP.eB vector solution was loaded into a 1 mL syringe using a 30-G needle without air bubbles. The tail was grasped at the distal end using the index and middle fingers while keeping the lower part of the tail held between the thumb and the ring fingers. The needle entered the vein at a shallow depth while maintaining the syringe and needle parallel to the tail. A total of 50 mL vector solution (1E+12 vg/mL, from the stock 2E+13 vg/mL) diluted in 100 mL sterile PBS was injected when a flash of blood was detected in the hub of the needle at the time of insertion without any resistance. After administration, the needle was removed, and gentle compression was applied until bleeding stopped. The animals recovered from the anesthesia were returned to their cage.

### Grip Strength Test

Grip strength analysis for both forelimbs and hind limbs was performed using a metal grid in a blind manner. Briefly the animals were placed on the center of a wire mesh cage top and inverted in such a way that only the forelimb and hindlimb paws could grasp the mesh. To perform the test, the grid was shaken gently enough for the animal to hold it and then turned upside down over an empty cage. The latency to fall (the duration each mouse was able to hang upside down) was measured in 3 trials for each animal with a maximum duration of 1 min. The mice were given ∼2 min interval between the attempts and the best attempt was used for the grip strength analysis.

### Survival Analysis

The body weight and overall health of all animals were monitored daily during the whole experiment to monitor the disease progression. Upon hindlimb paralysis, mice were provided wet chow and Dietgel. End stage was defined by the loss of the righting reflex, i.e., failure of mouse to right itself within 10 seconds when placed on its back. At this stage, the mice were euthanized, and tissue samples were collected for further analysis.

### Tissue Collection

Mice were euthanized and transcardial perfusion was done with ice-cold phosphate buffered saline (1X PBS, 100 mL) and 4% paraformaldehyde (PFA, 50mL, freshly prepared in 1X PBS). The brain and spinal cord (divided into cervical, thoracic, and lumbar regions) were dissected out and post-fixed overnight in ice-cold 4% PFA to process paraffin embedded sectioning. To extract the RNA, whole spinal cords were subjected to hydraulic extrusion using a 1 mL syringe and a 20-gauge needle after perfusion with 1X PBS. The tissues were flash frozen and stored at –80°C until further analysis.

### Histology and Immunohistochemistry

Formalin-fixed paraffin embedded sections were used for histology and immunohistochemistry analysis. 10 µm thick coronal sections of spinal cord were cut and stained with Cresyl violet (Nissl stain to label the total population of cells and show the cellular distribution in the area under investigation) for histological analysis. For immunohistochemistry, sections were de-paraffinized, subjected to antigen retrieval, and blocked at room temperature for 1 hour. The sections were then incubated overnight at 4°C with primary antibodies diluted in blocking buffer. The following primary antibodies were used, ChAT (Ref#AMAb91130, 1:1000, Atlas Antibodies), human-specific TDP-43 (Ref#WH0023435M1, 1:1000, Millipore Sigma) and mouse-specific TDP-43 (C-Terminus, Ref#12892-1-AP, 1:1000, Proteintech). CTR was detected using the human-specific TDP-43 antibody. Then, the sections were washed three times with 1X PBS containing 1% Tween-20 (PBST) followed by incubation in biotinylated secondary antibodies (Anti-Mouse IgG, Ref#BP-9200 and Anti-Rabbit IgG, Ref#BP-9100, Vector laboratories) diluted in blocking buffer at room temperature for 1 hour. Sections were washed three times with PBS-T and incubated with avidin-biotin peroxidase complex kit (Ref#PK-7100, Vectastatin ABC Reagent, Vector Laboratories) for 1 hour. After the incubation, the sections were washed three times with PBST and incubated with DAB solution for 30 seconds to 1 minute to produce the brown reaction product. The sections were counterstained with hematoxylin, dehydrated and mounted. The images were acquired using Zeiss Apotome Inverted Brightfield Microscope (Zeiss, Germany) and quantification of motor neurons was done using ImageJ software (National Institutes of Health, Bethesda, MD) from four serial sections per animal in a blinded manner independently by three investigators. The average number of total motor neurons was determined as normalized values to the total number of sections used.

### *In Situ* Hybridization: BaseScope^TM^ RED and Co-detection Assay

RNA *in situ* hybridization was performed using BaseScope Detection Reagent v2-RED Assay Kit (Ref#323900, Advanced Cell Diagnostics, ACDBio, USA) following the manufacturer’s instructions. 3ZZ probes were designed targeting the cryptic exon sequences within mouse *Synj2bp* (BA-Mm-Synj2bp-E2-intron2-NJ, Ref#712191, ACDBio, USA). To detect the infectivity of AAV, we have designed antisense probes against the vector backbone and performed BaseScope assay on spinal cord and brain tissue sections (BA-CMV-Enhancer-2zz-st-sense-C1, REF#1039721-C1, ACDBio, USA). To determine the accumulation of CTR transcripts, we have also designed a 1zz BaseScope probe against the CTR mRNA (BA-Hs-TARDBP-RAVER1-1zz-C1, Ref#1573691-C1, ACDBio, USA). Briefly, formalin-fixed paraffin-embedded sections (10 µm) were deparaffinized and pre-treated with hydrogen peroxide for 10 min at room temperature, target retrieval buffer for 15 min at 99°C and protease IV for 30 min at 40°C. The following day, sections were hybridized with the target probes in a HybEZ hybridization oven (ACDBio, USA) for 2 hours at 40°C. The hybridized signals were amplified through a series of amplification at 40°C and washing steps at room temperature. Finally, the signals were detected using a chromogenic red substrate and counterstained with hematoxylin. To assess the quality of RNA, each sample was hybridized with a probe targeting a mouse housekeeping gene (BA-Mm-Ppib, Ref#701071, ACDBio, USA). As a negative control to control for the background staining, each section was also evaluated with another probe targeting a bacterial gene (BA-DapB, Ref#701011, ACDBio, USA). The co-detection assay was performed using the RNA-Protein Co-Detection Ancillary Kit (Ref#323180, ACDBio, USA). The images were captured using a Zeiss Apotome Inverted Brightfield Microscope (Zeiss, Germany). The hybridized signal for each RNA transcript was detected as red punctate dots or clusters and quantified using ImageJ software (National Institutes of Health, Bethesda, MD).

### RNA extraction and RT-PCR

The tissues were homogenized using a Kimble Pellet Pestle Cordless Motor (Ref#749540, DWK Life Sciences) and total mRNA was extracted using RNeasy Mini kit (Ref#74106, Qiagen) following the manufacturer’s instructions. The extracted RNA was quantified using Nanodrop One (Ref#269-309101, Thermo Fisher Scientific, Inc.) and then converted to cDNA using the Protoscript II First Strand cDNA Synthesis Kit (Ref#E6550L, New England Biolabs Inc.). To detect the cryptic exon containing mRNA, we PCR amplified cDNA using primers specifically designed to target the cryptic exon and another primer spanning the canonical exon. We used primers against *Ift81*, forward: AAGTGCGAGGACTTCGTGAG, reverse: CAGCGATCTGTCTGCTTTGC and *Gapdh,* forward: AGGTCGGTGTGAACGGATTTG, reverse: GGGGTCGTTGATGGCAACA. *Gapdh* measurements were used for normalization. The resulting PCR products were then resolved on a 1.5% agarose gel containing ethidium bromide and visualized under UV light to detect the presence of a product at 187 bp, indicative of cryptic exon-containing mRNA.

### Ventral Root Extraction and Light Microscopy Analysis

To extract the lumbar dorsal root ganglion (DRG) with their corresponding roots, animals were perfused with freshly prepared 2% glutaraldehyde, 2% paraformaldehyde (EM grade prill), 50 mM sodium cacodylate, 50 mM phosphate (Sorenson’s) 3 mM MgCl_2_, pH 7.2 at 1085 mosmols. After perfusion, animals were kept in the cold room (4°C) for two hours and then the DRGs with corresponding roots were extracted and incubated in fresh fixative overnight at 4°C. Samples were processed as follows: All incubations were carried out at 4°C until the 70% ethanol step, then incubated at room temperature. Initially, samples were rinsed in a buffer consisting of 75 mM cacodylate 75 mM phosphate 3.5% sucrose 3 mM MgCl_2_, pH 7.2 with an osmolarity of 430 mOsm, for 45 minutes. Following the buffer rinse, samples were microwave-treated using a Pelco laboratory grade microwave model 3400, with a power setting of 50% and a pulse duration of 10 seconds, followed by a 20-second pause and then another 10-second pulse. The samples were then post-fixed in a solution containing 2% osmium tetroxide, 1.6% potassium ferrocyanide, and the same buffer without sucrose, and were placed on ice in the dark for 2 hours. Samples were then rinsed in 100 mM maleate buffer with 3.5% sucrose pH 6.2. This was followed by En-bloc staining of the samples for 1 hour with filtered 2% uranyl acetate in maleate sucrose buffer, pH 6.2. Samples were then dehydrated through a graded series of ethanol to 100%, transferred through propylene oxide, and embedded in Eponate 12 (Pella). The samples were then cured at 60°C for 2 days.

Each spinal ganglion with roots was embedded in epoxy resin and sagittal sections were cut on Reichert Ultracut E Microtome with a Diatome Diamond knife (450) to get 1 μm semithin sections. Toluidine blue staining was performed on each section to visualize the myelinated nerve fibers and images were acquired using a Zeiss Apotome Inverted Brightfield Microscope (Zeiss, Germany). Three animals per group were analyzed to quantify the nerve fibers. The quantifications were performed independently by three investigators in a blinded manner to the genotype and treatment.

### Muscle Histology

The hind limb muscles were dissected out and their gross morphology and weight were recorded. Each muscle was fixed in 4% PFA overnight, and 10 μm paraffin embedded sections were cut. The sections were deparaffinized and stained with hematoxylin and eosin (H&E). Myofiber morphology and diameter were assessed using a Zeiss Apotome Brightfield microscope (Zeiss, Germany).

### Statistical Analysis

Data analysis was performed through the following statistical tests using the Graphpad Prism v.5. software (GraphPad Software Inc., San Diego, CA, USA). Histological data was analyzed using an unpaired, two-tailed Student’s t-test with Tukey’s multiple comparison test and one-way analysis of variance (ANOVA) to identify significant differences between groups. Behavioral data was analyzed using two-way ANOVA with Turkey’s multiple comparison test. The Kaplan-Meier survival data were analyzed using log-rank test. The p-values less than 0.05 considered significant.

## Acknowledgment

We thank Michael Delannoy and Aswin Chandrasekhar for their technical assistance. This work was supported in part by the NIH grants (UG3/UH3NS115608, R33NS115161 and R01NS095969 to P.C.W.) the Kissick Family Foundation Frontotemporal Dementia Grant Program (to J.P.L.), and the Department of Defense (AL230138 to P.C.W.).

## Author contributions

P.C.W. and A.P.M. conceptualized, designed and interpreted the study. A.P.M., P.C.W. and J.P.L. wrote the manuscript. A.P.M., J.G.Y, O.S., M.B., S.R. and T.M. performed experiments and/or analyzed the data. I.R.S. aided with figure design. All the authors reviewed and approved the final manuscript.

## SUPPLEMENTARY FIGURES

**Supplementary Fig. 1:**
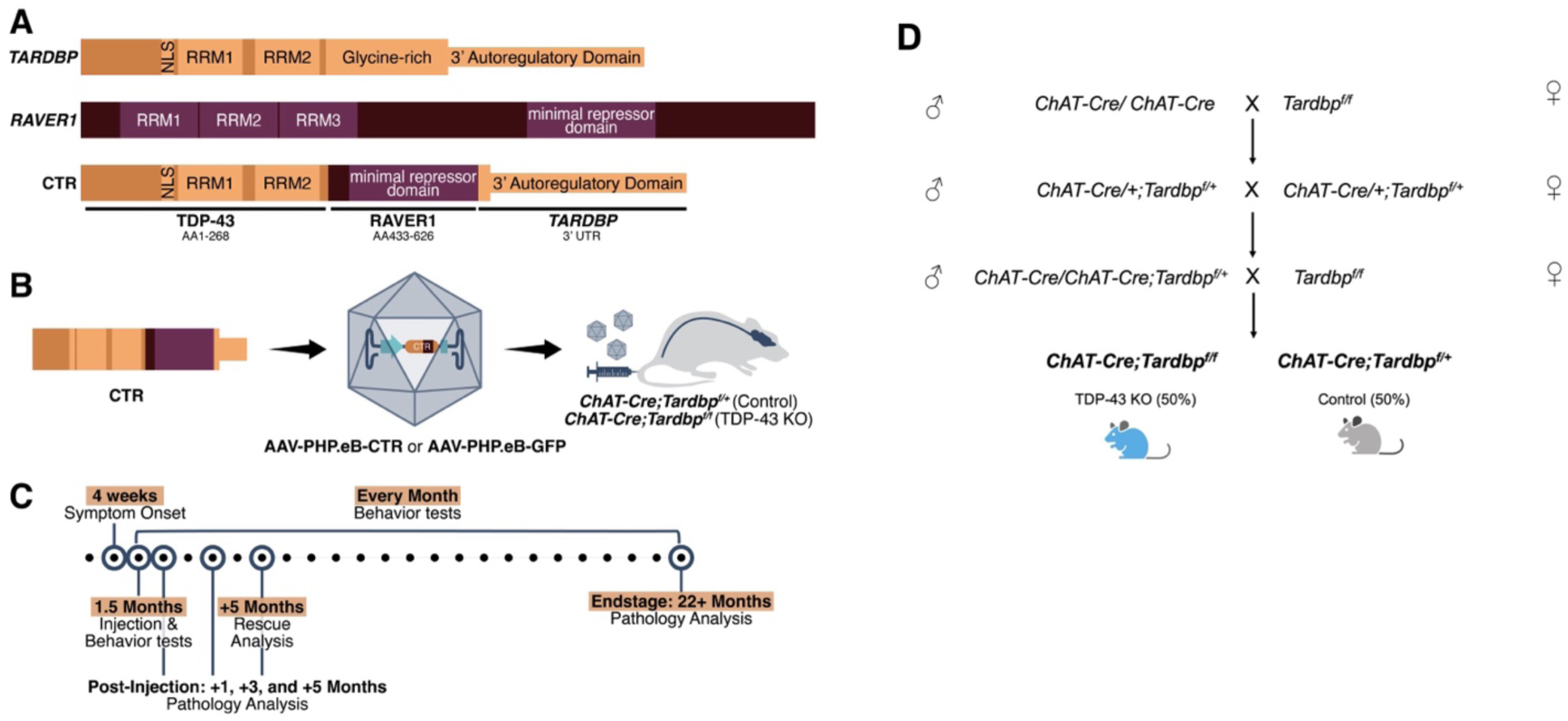
Schematic representations of structure of TDP-43, RAVER1, and CTR, CTR therapeutic strategy and study design. (**A**) The TDP-43 protein contains an N-terminal domain harboring a nuclear localization signal (NLS), two RNA-recognition motifs (RRMs), and a C-terminal domain which is glycine-rich and aggregation-prone. Parts of the final exon and 3’ untranslated region encode an autoregulatory domain. The RAVER1 protein contains an N-terminal domain with three RRMs and a C-terminal domain that functions as a splicing repressor. Our CTR is a fusion construct that contains N-terminal domain of *TARDBP* (encoding amino acid (AA) 1-268), with the NLS and both RRMs and the minimal repressor domain from *RAVER1* (encoding AA 433-626). The CTR also retains the 3’ autoregulatory domain from *TARDBP*. (**B**) The AAV-PHP.eB viral capsid, which can cross the blood-brain barrier, containing CTR transgene, is injected into *ChAT-Cre;Tardbp^f/+^* or *ChAT-Cre;Tardbp^f/f^* mice through tail vein injection. (**C**) Schematic of the temporal analysis showing behavior and molecular analyses. (**D**) Breeding strategy for generating TDP-43 knockout, *ChAT-Cre;Tardbp^f/f^* mice which shows TDP-43 depletion in ChAT positive motor neurons and littermate controls, *ChAT-Cre;Tardbp^f/+^* mice which maintain the normal level of TDP-43. % indicates the Mendelian frequency of pups from each litter expected to carry the genotype of interest.

**Supplementary Fig. 2:**
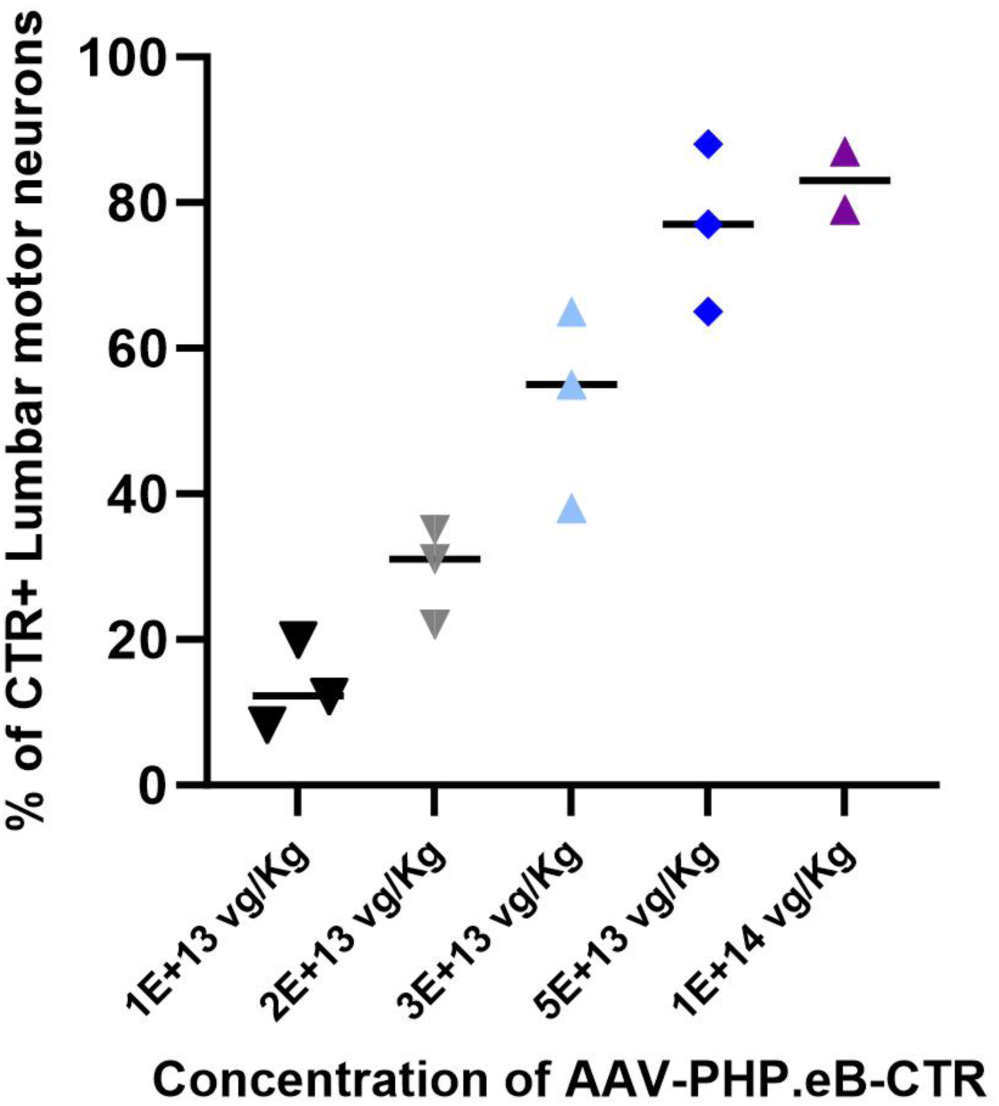
Optimization of the AAV-PHP.eB vector for intravenous delivery. The concentration of viral stock solution was 2E+13 vg/mL. Doses tested include 1E+13 vg/mL (n=3), 2E+13 vg/Kg (n=3), 3E+13 vg/Kg (n=3), 5E+13 vg/Kg (n=3) and 1E+14 vg/Kg (n=2). The optimum dose selected for the study was 5E+13 vg/Kg showing an average infectivity rate of 77%.

**Supplementary Fig. 3:**
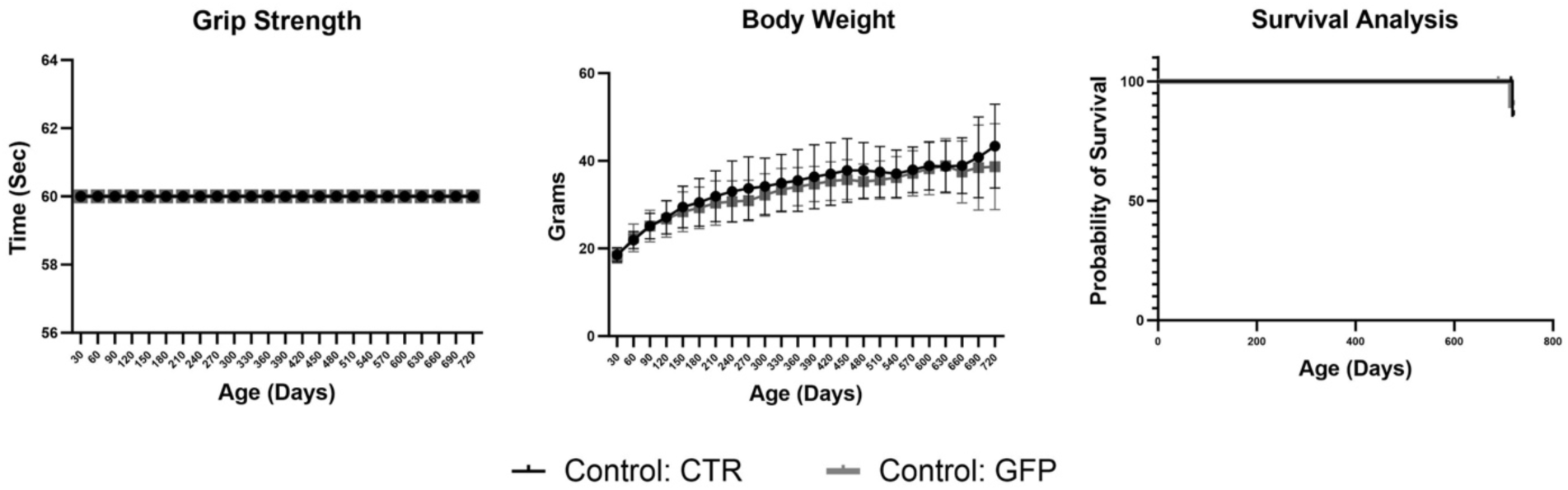
Evidence for no overt toxicity in aged *ChAT-Cre;Tardbp^f/+^* mice with long-term exposure to AAV-PHP.eB-CTR. The behavioral and survival analysis show no overt toxicity in *ChAT-Cre;Tardbp^f/+^* mice treated with AAV-PHP.eB-CTR suggesting the safety and efficacy of CTR (n=15).

**Supplementary Fig. 4:**
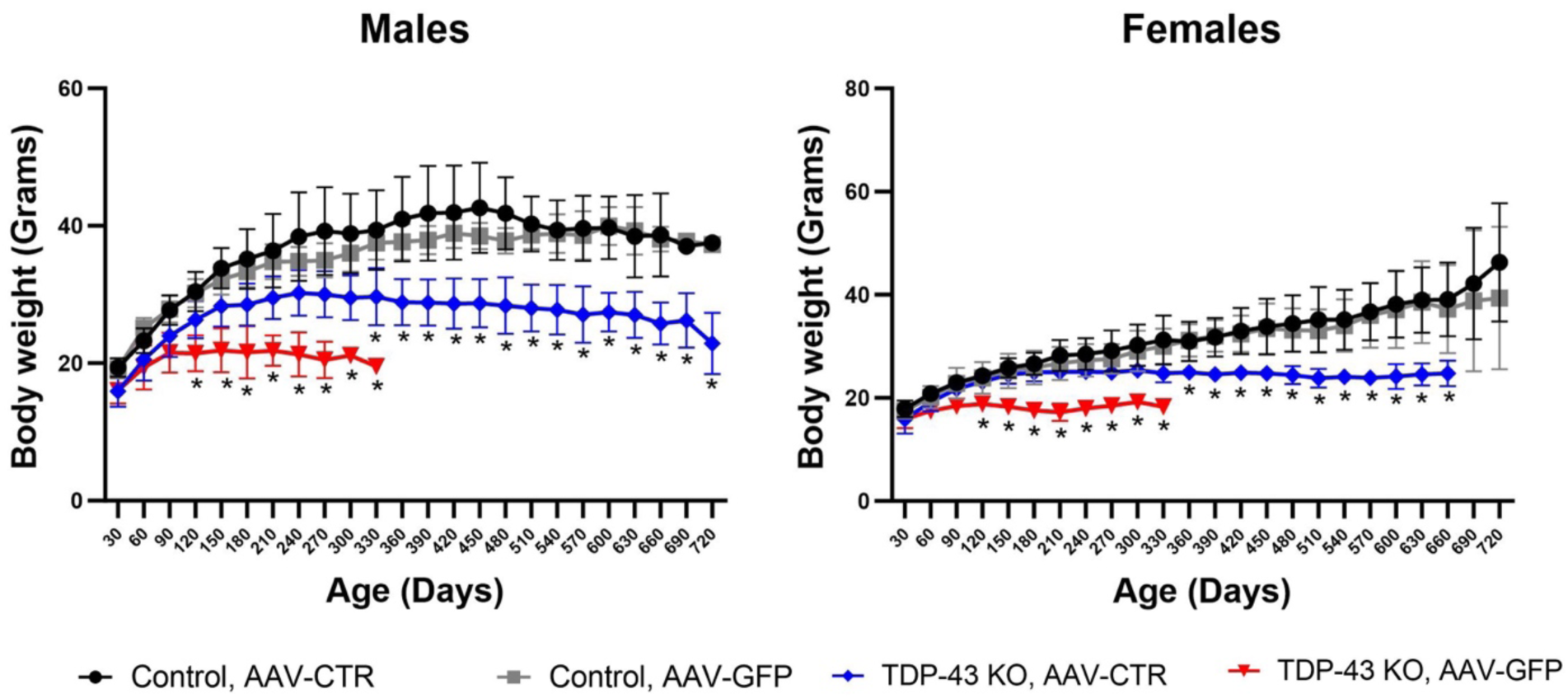
Body weight analysis in male (n=8) and female (n=8) cohort. Separate body weight analysis also showed significant improvement in the age-dependent reduction of body weight associated with the *ChAT-Cre;Tardbp^f/f^* mice (*p<0.0001, two way ANOVA using Turkey’s multiple comparison test).

**Supplementary Fig. 5:**
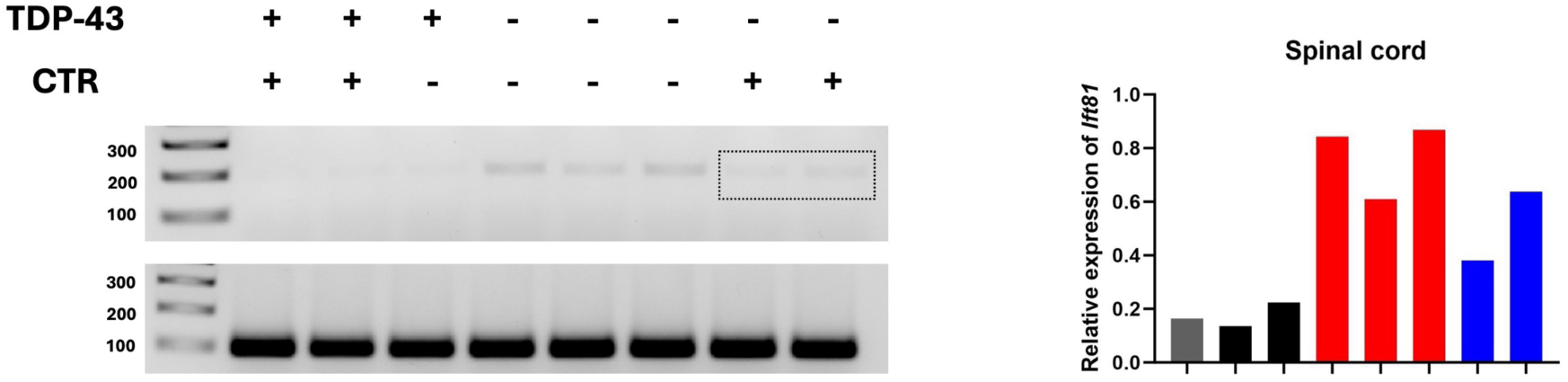
Reverse transcriptase-PCR analysis showing rescue of cryptic exon expression in *ChAT-Cre;Tardbp^f/f^* mice treated with AAV-PHP.eB-CTR. RT-PCR analysis using primers against *Ift81* cryptic exon in spinal cord shows that aberrant splicing of *Ift81* is rescued in *ChAT-Cre;Tardbp^f/f^* mice (n=2) treated with AAV-PHP.eB-CTR compared to the *ChAT-Cre;Tardbp^f/f^* mice (n=3) treated with AAV-PHP.eB-GFP.

**Supplementary Fig. 6:**
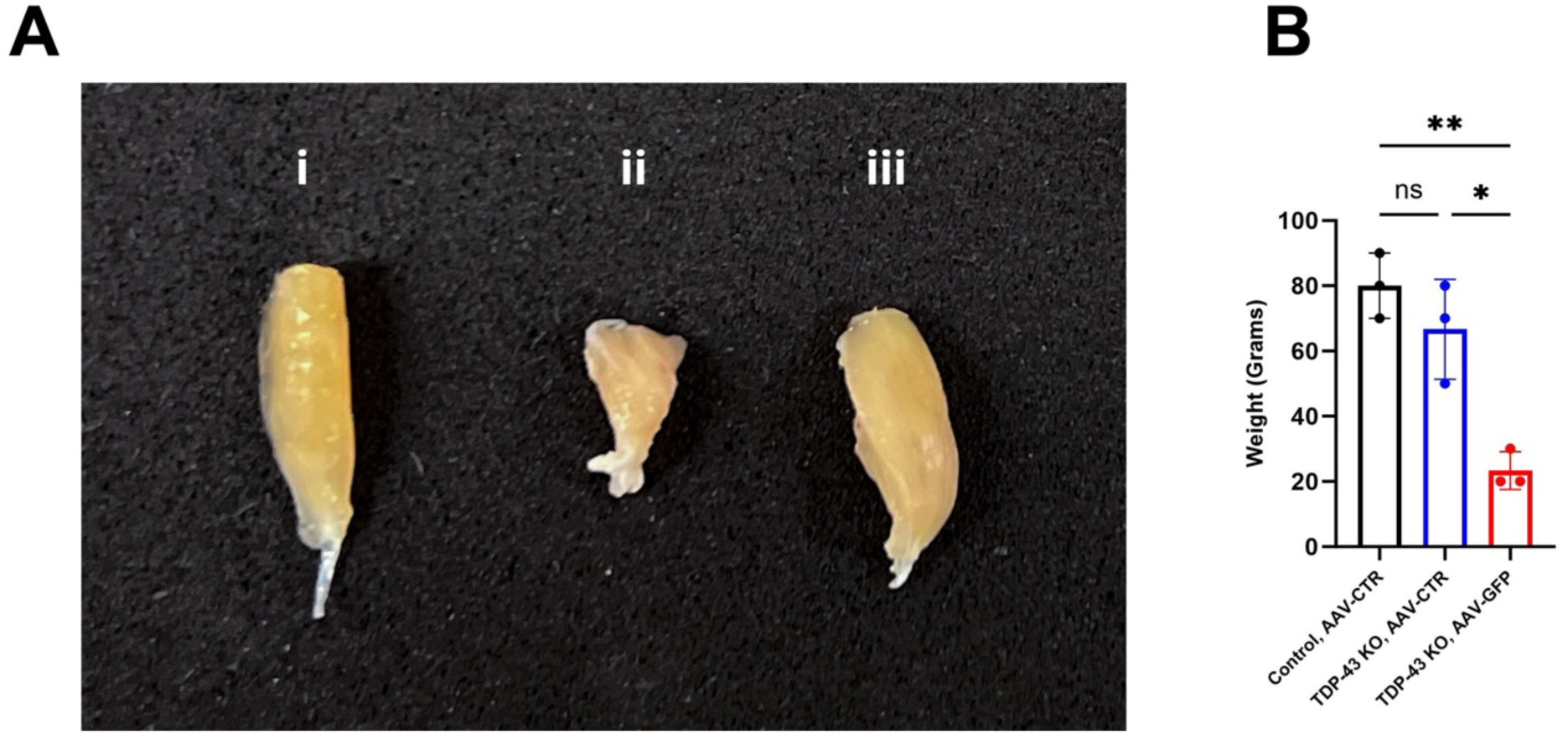
Gross analysis of hindlimb muscle size and weight shows rescue in *ChAT-Cre;Tardbp^f/f^* mice treated with AAV-PHP.eB-CTR. (**A**) Representative image of the gross structure of gastrocnemius muscle in *ChAT-Cre;Tardbp^f/+^* mice treated with AAV-PHP.eB-CTR (i), *ChAT-Cre;Tardbp^f/f^* mice treated with AAV-PHP.eB-GFP (ii) and *ChAT-Cre;Tardbp^f/f^* mice treated with AAV-PHP.eB-CTR (iii). (**B**) Average weight of the gastrocnemius muscle was significantly reduced in *ChAT-Cre;Tardbp^f/f^* mice treated with AAV-PHP.eB-GFP compared to *ChAT-Cre;Tardbp^f/+^*(p value=0.006) and is rescued in *ChAT-Cre;Tardbp^f/f^* mice treated with AAV-PHP.eB-CTR compared to *ChAT-Cre;Tardbp^f/+^* treated with AAV-PHP.eB-CTR (p value=0.44) and *ChAT-Cre;Tardbp^f/f^* mice treated with AAV-PHP.eB-GFP (p=0.006) (n=3 per group).

